# Role of glutamate 292 and lysine 331 in catalysis for the flavoenzyme (S)-6-hydroxynicotine oxidase from *Shinella* sp. HZN7

**DOI:** 10.1101/2025.05.21.655333

**Authors:** Zhiyao Zhang, Kaitrin Freeland, Frederick Stull

## Abstract

The flavoenzyme NctB from *Shinella* sp. HZN7 catalyzes the oxidation of (S)-6-hydroxynicotine to 6-hydroxypseudooxynicotine concomitant with dioxygen reduction, which is the same chemistry catalyzed by the well-studied flavoenzyme L-6-hydroxynicotine oxidase (LHNO) from *Paenarthrobacter nicotinovorans*. However, while both enzymes are members of the flavoprotein amine oxidoreductase (FAO) family, they share only 26% sequence identity and are evolutionarily distant. Furthermore, nearly all FAOs (including LHNO) have a conserved lysine proximal to N5 of the flavin that is known to promote the reaction with O_2_ in the oxidative half-reaction, yet NctB, unusually, has a glutamate (Glu292) at this position. We report here using transient kinetics that NctB reacts rapidly with dioxygen in the oxidative half-reaction despite lacking the conserved lysine associated with promoting the reaction with O_2_ in FAO family enzymes. Mutagenesis reveals that a lysine derived from a different sequence position (Lys331) likely accelerates the reaction with dioxygen in NctB, as the K331M mutation results in a 1400-fold decrease in rate constant for reaction with O_2_. Glu292 forms a salt bridge with Lys331 in the structure of NctB, and a E292T mutation results in a ∼80-fold decrease in rate constant for reaction with O_2_, suggesting that Glu292 optimizes the positioning and/or properties of Lys331 to promote dioxygen activation. Analysis of pH-rate effects in NctB shows similar pH profiles as in LHNO despite having differences in active site structure. These results indicate that NctB and LHNO convergently evolved to have the same enzymatic function.

## INTRODUCTION

Several groups of bacteria have been identified that are capable of catabolizing nicotine for use as a carbon and nitrogen source [1–4]. Three different pathways have been identified that use different sets of enzymes for nicotine catabolism: the pyrrolidine pathway, the pyridine pathway, and the variation of the pyridine and pyrollidine (VPP) pathway, which is a hybrid of the other two pathways [1,3,5]. The pyrrolidine pathway begins with the enzyme nicotine oxidoreductase (NicA2), which oxidizes a carbon-nitrogen bond in the pyrrolidine ring of nicotine to yield N-methylmyosmine, which spontaneously hydrolyzes into pseudooxynicotine (Scheme 1) [4,6]. The pyridine pathway instead starts with hydroxylation of the pyridyl ring of nicotine to yield (S)-6-hydroxynicotine (6OHN), catalyzed by nicotine dehydrogenase. Then a carbon-nitrogen bond in the pyrrolidine ring of 6OHN is oxidized by L-6-hydroxynicotine oxidase (LHNO) to yield 6-hydroxy-N-methylmyosmine, which hydrolyzes into 6-hydroxypseudooxynicotine (6HPON) (Scheme 1) [3,7]. The VPP pathway is a hybrid of the pyrrolidine and pyridine pathway where the first two enzyme-catalyzed steps in the VPP pathway are the same as the pyridine pathway, but 6HPON is eventually converted into 6-hydroxy-3-succinoyl-pyridine and joins the pyrrolidine pathway thereafter [3,8–10].

LHNO and NicA2 are related enzymes that are members of the flavoprotein amine oxidoreductase (FAO) superfamily. These enzymes use a flavin adenine dinucleotide (FAD) prosthetic group to perform a two-electron oxidization of a C–N bond in their substrate in the reductive half reaction, reducing the FAD to the hydroquinone form (FADH_2_) in the process (Scheme 2). The FADH_2_ is then reoxidized to FAD in the oxidative half reaction by a second substrate molecule, which is O_2_ for most FAOs, including LHNO, but is a cytochrome c protein for NicA2 [11–13]. A sequence and structural feature that is present in almost all FAOs is a lysine residue that forms a water-mediated hydrogen bond with N5 of the flavin. This highly conserved lysine residue has been shown to be critical for enhancing the rate of reaction with O_2_ in several oxidases of the FAO superfamily [14–21]. The lysine is thought to provide a positive charge that promotes the transient formation of superoxide and flavin semiquinone, which is a necessary step *en route* to forming H_2_O_2_ and oxidized flavin, and the lysine is also thought to serve as a proton donor during O_2_ reduction by FADH_2_ [14].

Prior phylogenetic analyses have shown that the LHNO enzymes associated with the VPP pathway (VPP-LHNO) are distantly related to the LHNO enzymes from the pyridine pathway (Pyridine-LHNO) [1]. The VPP-LHNO enzymes are more closely related to NicA2-like enzymes, with the VPP-LHNO and NicA2 groups recently diverging from a common ancestor, whereas the Pyridine-LHNO enzymes are present in a completely separate lineage from that of NicA2 and VPP-LHNO [22]. As a further illustration of this point, the VPP-LHNO from *Shinella* sp. HZN7 (NctB) shares 43% sequence identity with NicA2 from *Pseudomonas putida* S16, but only shares 26% sequence identity with *Paenarthrobacter nicotinovorans* Pyridine-LHNO. Thus, phylogenetic analysis suggests that VPP-LHNO enzymes and Pyridine-LHNO enzymes independently evolved to catalyze the same reaction [22]. Curiously, the VPP-LHNO enzymes all have a glutamate (Glu292 in NctB) at the homologous sequence position of the lysine that is highly conserved among FAOs (Fig. 1A). However, several VPP-LHNO enzymes have been biochemically characterized and shown to react rapidly with O_2_ [3,23–25], indicating that replacement of the highly conserved lysine with glutamate has not compromised their ability to react quicky with O_2_. The recently solved structure of NctB shows that a lysine at a different sequence position (Lys331) has its side chain oriented in front of the flavin N5, leading us to hypothesize that this alternative lysine in VPP-LHNO enzymes plays a similar role as the lysine that is conserved in *P. nicotinovorans* Pyridine-LHNO (Lys287) and most other FAO family enzymes (Fig. 1A-1C) [11]. Our prior studies on ancestral sequence reconstructions indicate that the change in lysine sequence position occurred when the enzymes ancestrally had dehydrogenase function and initially involved replacement of the highly conserved lysine with a threonine (Fig. 2) [22]. The threonine was later replaced with a glutamate (Glu292 in NctB) when the ancestral enzymes gained oxidase function, and the glutamate is retained at this position in all modern day VPP-LHNO enzymes (Fig. 2) [22]. In the structure of NctB, Glu292 is positioned to form a favorable charge-charge interaction with Lys331 (Fig. 1B), which may affect the properties and/or positioning of Lys331 in the active site. We describe here the use of site-directed mutagenesis in NctB to interrogate the role of this alternative lysine architecture in catalysis and O_2_ activation for VPP-LHNO enzymes.

**Fig. 1.**
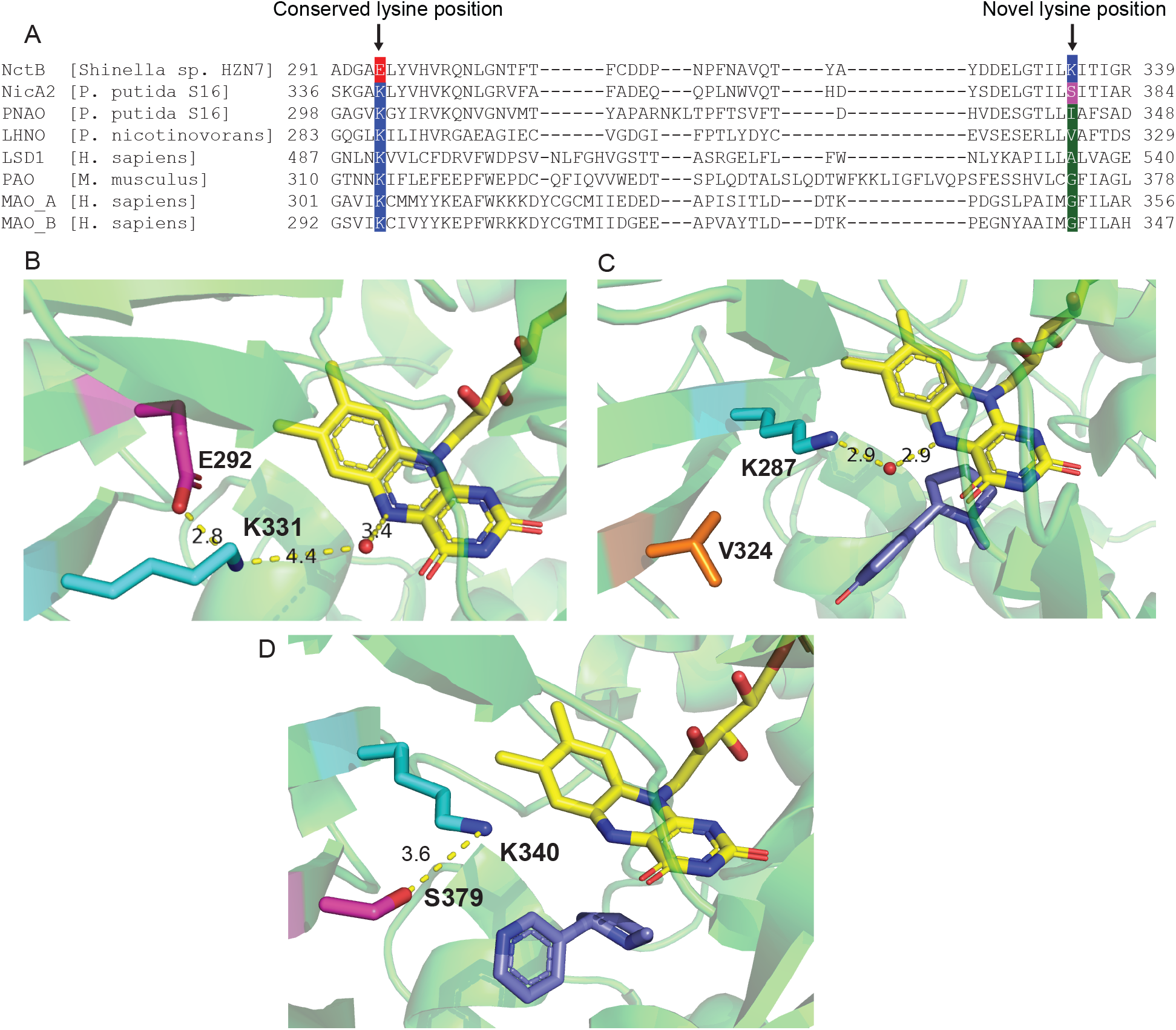
Alignment and active sites of NctB and related enzymes. (A) Sequence alignment of NctB and other FAO family enzymes that contain the conserved lysine. PNAO is pseudooxynicotine amine oxidase [36]; LSD1 is lysine-specific demethylase 1 from *H. sapiens* [39]; PAO: N1-acetylpolyamine oxidase from *M. musculus* [40]; MAO A/B are monoamine oxidase A/B from *H. sapiens* [41,42]; (B) Active site of NctB (PDB 6CR0) depicting Glu292, the novel lysine (Lys331) and a crystallographic water molecule. (C) Active site of Pyridine-LHNO from *P. nicotinovorans* (PDB 3NG7) depicting the lysine at the highly conserved sequence position (Lys287) and the water-mediated hydrogen bond to N5 of the flavin. 6OHN is bound in this structure (shown in dark blue) (D) Active site of NicA2 (PDB 6C71). NicA2 backbone is shown in green. K340 is shown in blue and S379 is colored in magenta. Nicotine is bound in this structure (shown in dark blue).

**Fig. 2.**
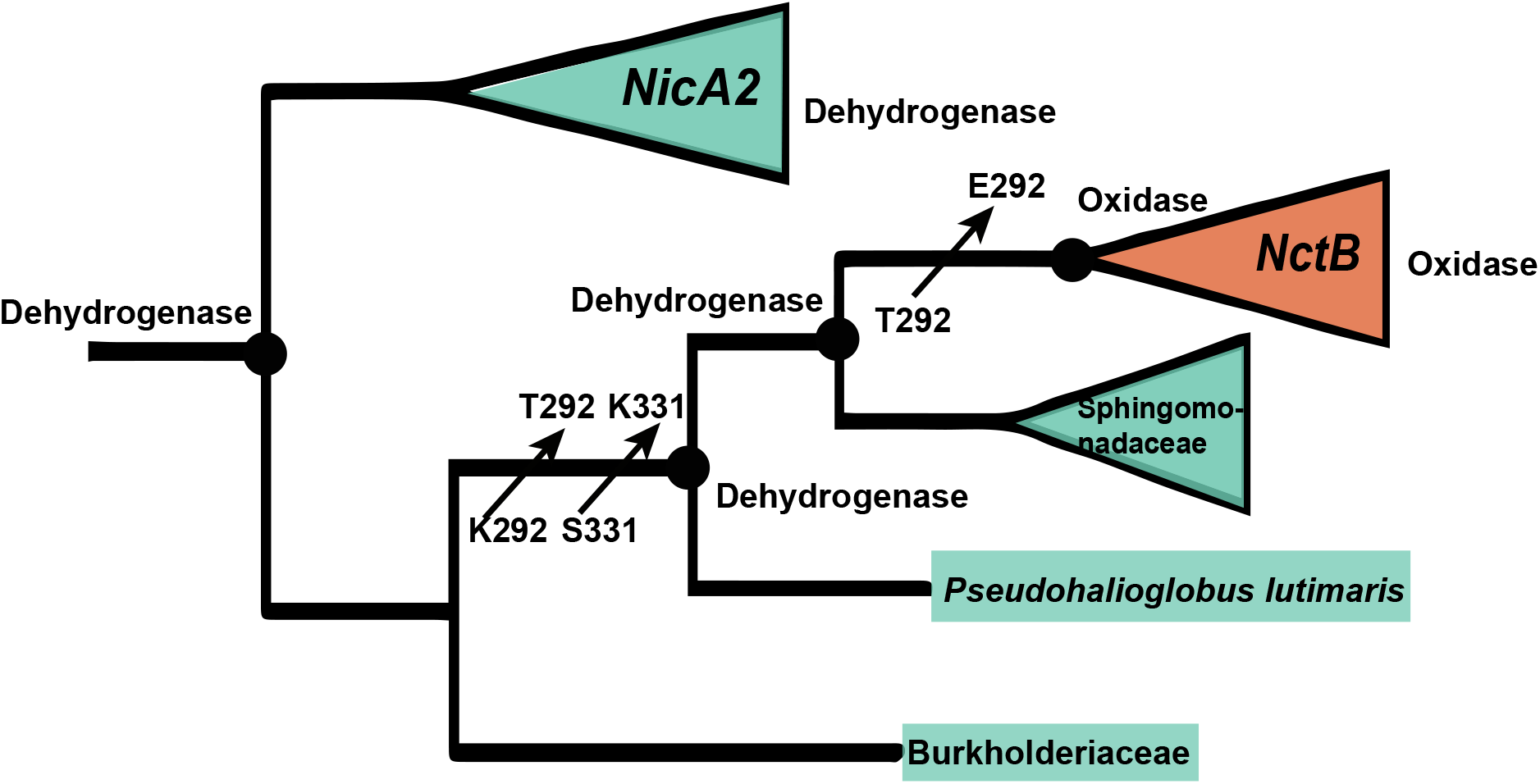
Phylogenetic tree of Nicotine/6OHN degrading enzymes. Collapsed phylogeny of nicotine/6OHN degrading flavoprotein amine oxidoreductases. Clades and individual enzymes are colored based on dehydrogenase (green) or oxidase (orange) function. The “Dehydrogenase” and “Oxidase” labels for ancestral nodes and modern-day enzymes refer to data collected in a prior study on ancestral sequence reconstructions [22]. The numbering for amino acid substitutions is based on the sequence of NctB from *Shinella* sp. HZN7.

## MATERIALS AND METHODS

### Materials

The 6-hydroxynicotine used in this study was acquired as a racemic mixture of the (R) and (S) isomers from Enamine (catalog # EN300-32297860), similar to that used previously for characterization of *P. nicotinovorans* Pyridine-LHNO [7,14]. 6OHN concentrations reported in this paper denote the concentration of the (S) stereoisomer, which is the substrate for LHNO enzymes. The plasmid used for expression of NctB was obtained from Twist Biosciences as a synthetic construct cloned into the NdeI and XhoI sites of pET29b to generate a construct with a C-terminal His tag. Mutants of NctB were prepared via site-directed mutagenesis using PCR and mutagenic primers via the Quickchange method. The introduction of mutations was verified by Sanger sequencing.

### Protein expression and purification

The WT NctB (or mutant) expression plasmid was introduced into *E. coli* BL21 (DE3) cells and cultivated in expression media consisting of 12 g/L tryptone, 24 g/L yeast extract, 40 ml/L glycerol, 0.072M K_2_HPO_4_, and 0.017M KH_2_PO_4_ (PEM). Cultures were incubated at 37 °C with shaking until an OD_600_ of 0.8. The temperature was lowered to 20 ^o^C, and cultures were induced with 100 μM IPTG and left to grow overnight. Pellets were harvested and resuspended in lysis buffer (300 mM NaCl, 50 mM NaH_2_PO_4_, 20 mM imidazole, 10% v/v glycerol, pH 8). The resuspended cells were lysed by sonication after EDTA-free protease inhibitor cocktail (Abcam) and Benzonse nuclease (Sigma) were added. The lysate was cleared by centrifugation at 24000 x g for 25 minutes. Supernatant was collected and loaded onto a nickel affinity column (resin from GBiosciences) that had been pre-equilibrated with lysis buffer. The column then was washed with 10 column volumes (CV) of lysis buffer and proteins were eluted with 2 CV of elution buffer (300 mM NaCl, 50 mM NaH_2_PO_4_, 250 mM imidazole, 10% v/v glycerol, pH 8). Proteins were concentrated and then subjected to an additional purification by passing through a HiLoad^TM^ 16/600 Superdex^TM^ 200 pg column at 4 ^O^C. This purification process was carried out using buffer consisting of 40 mM Hepes, 100 mM NaCl, and 10% v/v glycerol, at pH 7.5 (stopped-flow buffer). The purified enzymes were concentrated and flash frozen for storage at - 80×C. The extinction coefficient of NctB in stopped-flow buffer was determined by denaturing the enzyme with 1% SDS and 95 °C treatment for 5 minutes to release free FAD. The extinction coefficient of FAD bound to NctB was then calculated from the predenaturation and postdenaturation spectra using the known ε_450_ of 11,300 M^-1^cm^-1^ for free FAD. Using this method, an ε458 value of 11,600 M^−1^cm^−1^ was determined for NctB.

### Transient kinetic assays

All transient kinetics experiments were done in stopped-flow buffer at 4 ^o^C using a TgK Scientific SF-61DX2 KinetAsyst stopped-flow instrument. All protein solutions were made anaerobic in glass tonometers by repeated cycles with vacuum and anaerobic argon [26]. When needed for oxidative half-reaction experiments, the FAD group in NctB (or variant) was reduced in the anaerobic tonometer by titrating with 10 mM anaerobic 6OHN in a gas tight hamilton syringe. Anaerobic 6OHN was prepared by sparging for at least 10 min with anaerobic argon. The anaerobic 6OHN was slowly added to the point where the FAD group was fully reduced to the hydroquinone state, with the titration process monitored spectrophotometrically using a Cary 50 Bio UV–Visible Spectrophotometer and Cary WinUV software. For experiments monitoring the reaction with O_2_, NctB containing FAD hydroquinone in a tonometer was loaded on the stopped-flow instrument and mixed with buffer containing various oxygen concentrations (prepared by sparging buffer with various N_2_/O_2_ ratios made using a gas blender). The O_2_ concentration for each N_2_/O_2_ ratio was measured using a Hansatech Oxygraph+ oxygen probe system prior to loading on the stopped-flow. For experiments monitoring the reaction with 6OHN, oxidized anaerobic NctB was mixed with various concentrations of 6OHN (made anaerobic by sparging with argon), and the reaction was monitored using the instrument’s single-wavelength photomultiplier tube detector at 450 nm.

Stopped-flow data were analyzed using Kaleidagraph. Kinetic traces at 450 nm for the reaction with oxygen for were fit to a single exponential function (Equation 1) to determine the observed rate constant (k_obs_) for each oxygen concentration. Kinetic traces for reaction with 6OHN in the reductive half-reaction were fit to a sum of two exponentials (Equation 2) to determine k_obs_ values for the two kinetic phases, except for E292T variant which shows an extra third phase (Equation 3). The kinetic amplitude for each phase is represented by ΔA, the apparent first-order rate constant is denoted as k_obs_, and A∞ refers to the absorbance at the completion of the reaction.

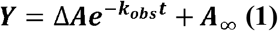

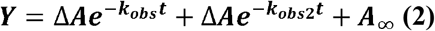

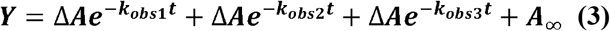

Plots of k_obs_ against substrate concentrations for reaction with oxygen were linearly fitted with the slope providing the second-order rate constant between the reaction of reduced enzyme and oxygen (k_ox_^O2^). Plots of k_obs_ against and 6OHN displayed a hyperbolic dependence and were fit to Equation 4 to determine k_red_, the rate constant for flavin reduction and K_d_ for substrate binding to oxidized enzyme.

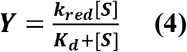

### pH dependent assays

Initial rates of amine oxidation were measured by monitoring the rate of oxygen consumption using a Hansatech Oxygraph+ oxygen probe system. All experiments were completed at 4 ^o^C. The buffers were 40 mM HEPES for pH 7-8, 40 mM CHES for pH 8.5-10, 40 mM MES at pH 5.5-6.5; all buffers also contained 0.1 M NaCl and 10% glycerol. Assays typically contained 200 nM enzyme with the exception of the lowest and highest pH, where 2 uM enzyme was used, and the reactions were initiated by added enzyme to the assay last. Apparent k_cat_ and k_cat_/K_m_ values for 6OHN were determined by varying concentration of the amine substrate at constant, ambient O_2_ concentration (∼400 μM at 4 ^o^C) and fitting the data to the Michaelis-Menten equation. Kaleidagraph was used to fit the pH value against the log of k_cat_, or k_cat_/K_m_ values. The pKa values for the k_cat_/K_m_ data were determined based on equation 5 below, where *c* is the maximum reaction rate and H is the concentration of hydrogen ion.

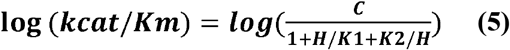

### Flavin Reduction Potential Measurements

Spectrophotometric measurements of NctB E_m_ values were performed on a Shimadzu UV-2501PC scanning spectrophotometer at 25 °C. Reduction potentials were determined by the xanthine/ xanthine oxidase method developed by V. Massey [27,28]. Briefly, a solution of 16 μM NctB (or mutants), 300 μM xanthine, 0.5 μM benzyl biologen, and anthroquinone-2,6-disulfonate at a concentration where A_375_∼0.2 as indicator dye (E_m_=-184 mV) in stopped flow buffer was made anaerobic in an anaerobic cuvette by repeated cycles with vacuum and anaerobic argon; a final concentration of 0.25 μM xanthine oxidase was also made anaerobic in the side arm of the anaerobic cuvette. The experiment was initiated by mixing in the xanthine oxidase from the side arm, and absorbance spectra from 700 to 250 nm were recorded every 10-15 min for ∼10-18 h. 550 nm was used to monitor the reduction of the dye since the FAD in NctB does not absorb at this wavelength. 450 nm was used to monitor reduction of NctB. However, reduced anthraquinone-2,6-disulfonate also absorbs at this wavelength, so data points at 450 nm were corrected to remove the contribution of the dye. Reduction of anthraquinone-2,6-disulfonate in the absence of NctB showed that reduced dye has an A450/A550 ratio of 5.2 whereas oxidized dye does not absorb at either of these two wavelengths. Since NctB does not contribute to the signal at 550 nm, the contribution of reduced anthraquinone-2,6-disulfonate at 450 nm could be determined by multiplying the signal at 550 nm by 5.2 for every individual scan. This value was then subtracted from the signal at 450 nm to isolate the signal of NctB at 450 nm for each individual scan. Data was fitted to the Nernst equation as previously described for the absorbance changes of the enzyme and dye using eq. 6. n is the number of electrons transferred (for 2 electron transfer), F is the Faraday constant, R is the gas constant, and T is the temperature in Kelvin.

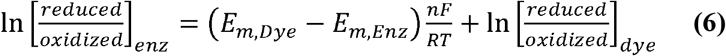

The midpoint potentials for enzyme and dye are E_m,Enz_ and E_m,Dye_ respectively. Subscripts for dye and enzyme refer to the fractional reduction values monitore for each (550 nm for dye, corrected 450 nm for the enzyme). The midpoint potentials were extracted from linear regression analysis of Nernst plots.

## RESULTS

### Rapid reaction kinetics

The kinetics of the reductive and oxidative half-reactions for NctB were independently measured by stopped-flow spectroscopy, using the changes in flavin absorbance at 450 nm as a readout (Fig. 3). The oxidative half-reaction (reaction between O_2_ and NctB containing FADH_2_) at 4°C showed a single-phase increase in absorbance at 450 nm, corresponding to flavin oxidation, and kinetic traces were fit to a single exponential function (eq 1) (Fig. 3A). The observed rate constant (k_obs_) for the reaction varied linearly with O_2_ concentration, consistent with a second order reaction, as is typically observed for most flavoproteins (Fig. 3A, inset) [29]. Linear fitting of the k_obs_ vs [O_2_] plot provided the second order rate constant for flavin oxidation by O_2_ (k_ox_^O2^) of 3.8±0.1*10^4^ M^-1^s^-1^ for NctB (Table 1). This value is comparable to that previously reported for *P. nicotinovorans* Pyridine-LHNO (2.7 ± 0.6*10^5^ M^-1^s^-1^), indicating that NctB reacts rapidly with O_2_ despite lacking the lysine at the sequence position that is typically conserved among FAO family enzymes [7].

**Table 1.**
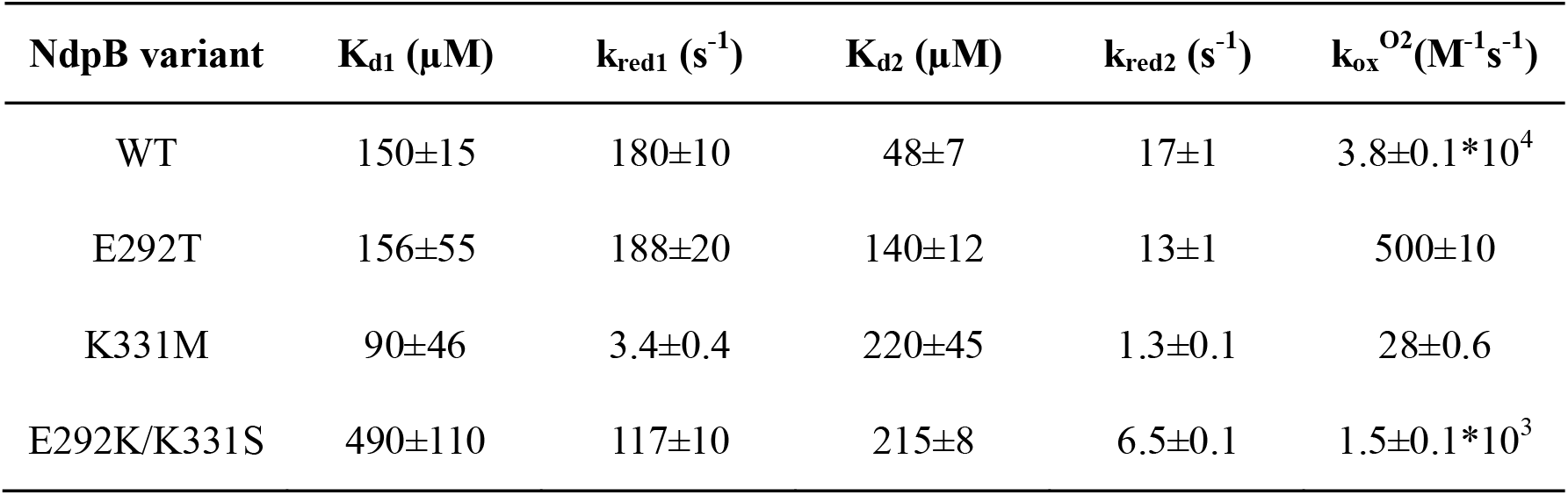
Kinetic Parameters of NctB and mutants

| **NdpB variant** | **K<sub>d1</sub> (μM)** | **k<sub>red1</sub> (s<sup>-1</sup>)** | **K<sub>d2</sub> (μM)** | **k<sub>red2</sub> (s<sup>-1</sup>)** | **k<sub>ox</sub><sup>O2</sup>(M<sup>-1</sup>s<sup>-1</sup>)** |
| --- | --- | --- | --- | --- | --- |
| WT | 150±15 | 180±10 | 48±7 | 17±1 | 3.8±0.1*10<sup>4</sup> |
| E292T | 156±55 | 188±20 | 140±12 | 13±1 | 500±10 |
| K331M | 90±46 | 3.4±0.4 | 220±45 | 1.3±0.1 | 28±0.6 |
| E292K/K331S | 490±110 | 117±10 | 215±8 | 6.5±0.1 | 1.5±0.1*10<sup>3</sup> |

**Fig. 3.**
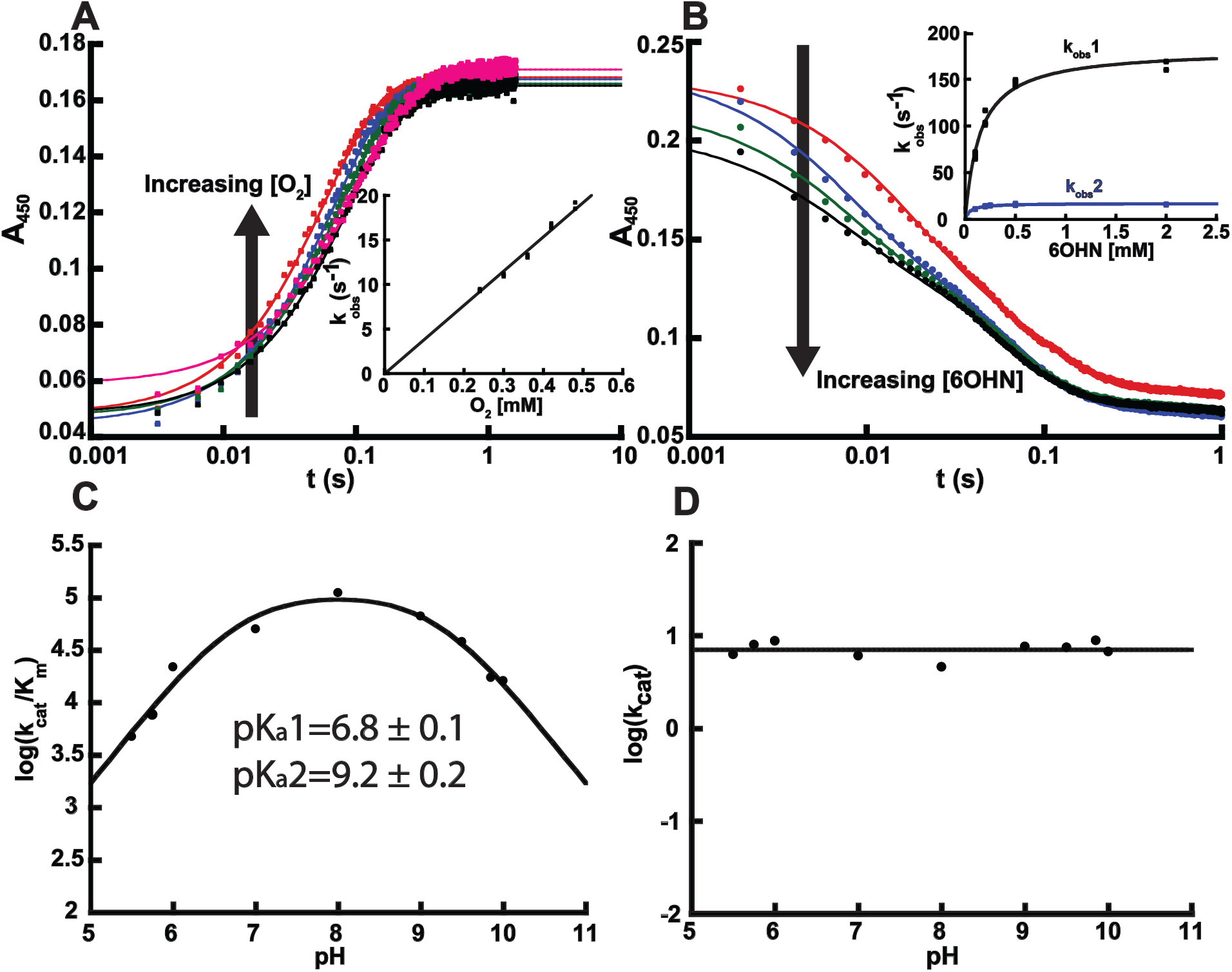
Half-reactions and pH profile for wild-type NctB. (A) Stopped-flow absorbance traces at 450 nm for the oxidation of reduced NctB by various concentrations of O_2_. Note the logarithmic time scale. The inset shows the plot of k_obs_ values against O_2_ concentration for the reaction of reduced NctB with O_2_. The slope obtained from linear fitting gives the k_ox_^O2^ value for NctB reported in Table 1. (B) Absorbance trace overlay at 450 nm for the reduction of NctB by various concentrations of 6OHN. Note the logarithmic time scale. The inset shows the k_obs_ values for the first (black) and second (blue) phase plotted against the concentrations of 6OHN. Both phases showed hyperbolic dependence and were fitted to Equation 4. The k_red_ and K_d_ values associated with these data sets can be found in Table 1. (C) pH dependence of log(k_cat_/K_m_). Fitting the data to eq.5 yielded apparent pKa values of 6.8 ± 0.1 and 9.2 ± 0.2. (D) pH dependence of log(k_cat_).

Stopped-flow traces for the reductive half reaction (reaction between 6OHN and NctB containing oxidized FAD) at 4°C display a biphasic decrease at 450 nm, with the reaction completing in one second (Fig. 3B). Kinetic traces fit best to a sum of two exponentials (eq 2), with the two phases having similar kinetic amplitudes. Similar biphasic behavior with comparable amplitudes was observed previously in the reaction between NicA2 and nicotine, and this was interpreted as the two active sites of the enzyme homodimer reacting with amine substrate with different kinetics [12,30]. The k_obs_ for both kinetic phases varied with 6OHN concentration, with both plateauing at a limiting value at high 6OHN concentrations (Fig. 3B, inset). Plots of k_obs_ versus [6OHN] were therefore fit to eq 4 to determine the first-order rate constant for flavin reduction when 6OHN is bound (k_red_) and the apparent K_d_ value for binding of 6OHN to oxidized NctB. The obtained kinetic parameters are listed in Table 1.

### pH dependence of steady state kinetic parameters

Steady state kinetic parameters for the NctB were determined at 4°C over the pH range of 5.5-10 by varying the 6OHN concentration at ambient concentrations of O_2_ (∼400 μM at 4°C); the enzyme was not stable outside of this pH range. The resulting kinetic parameters should be treated as apparent values since they were only collected at a single, fixed O_2_ concentration. As shown in Fig. 3C, a plot of log(k_cat_/K_m)_ against pH displayed a bell-shape profile and therefore was fit with eq. 5 to determine pKa values of 6.8±0.1 and 9.2±0.2 for the acidic and basic limbs of the profile, respectively. The k_cat_ versus pH profile was constant across the pH range tested (Fig. 3D).

### NctB’s alternative lysine architecture is critical for its rapid reaction with O_2_

Transient kinetic parameters for the reductive and oxidative half reactions were measured for Glu292Thr and Lys331Met variants of NctB, as these two residues constitute the unusual, alternative lysine architecture proximal to N5 of the flavin in NctB and VPP-LHNO enzymes (Fig. 1). Threonine was chosen as the replacement for Glu292 because a Thr to Glu substitution occurred at this position when oxidase function evolved in the VPP-LHNO lineage from a dehydrogenase ancestor that reacts poorly with O_2_ (Fig. 2) [22]. Methionine was chosen as the replacement for Lys331 since methionine is isosteric with lysine but lacks the amine on its side chain. The Lys331Met variant had a k_ox_^O2^ of 28 M^-1^s^-1^, ∼1400 fold lower than that of WT, confirming that the lysine at the alternative sequence position in NctB is critical for the enzyme’s ability to react rapidly with O_2_ (Fig. 4A, Fig. 4D and Table 1). The k_red_ values for Lys331Met variant were also 53 and 13-fold lower than in WT, indicating that this residue plays a role in the reaction with 6OHN in addition to its contribution to the oxidative half reaction (Fig. 5A, 5D and Table 1). The Glu292Thr variant displayed a ∼80-fold decrease in k_ox_^O2^ compared with wild type, highlighting the importance of Glu292 in oxygen activation by NctB (Fig. 4B,4D and Table 1). In contrast, the reductive half reaction was unaffected by the Glu292Thr mutation, as the k_red_ values for this variant were indistinguishable from wild type (Fig. 5B, 5E and Table 1).

**Fig. 4.**
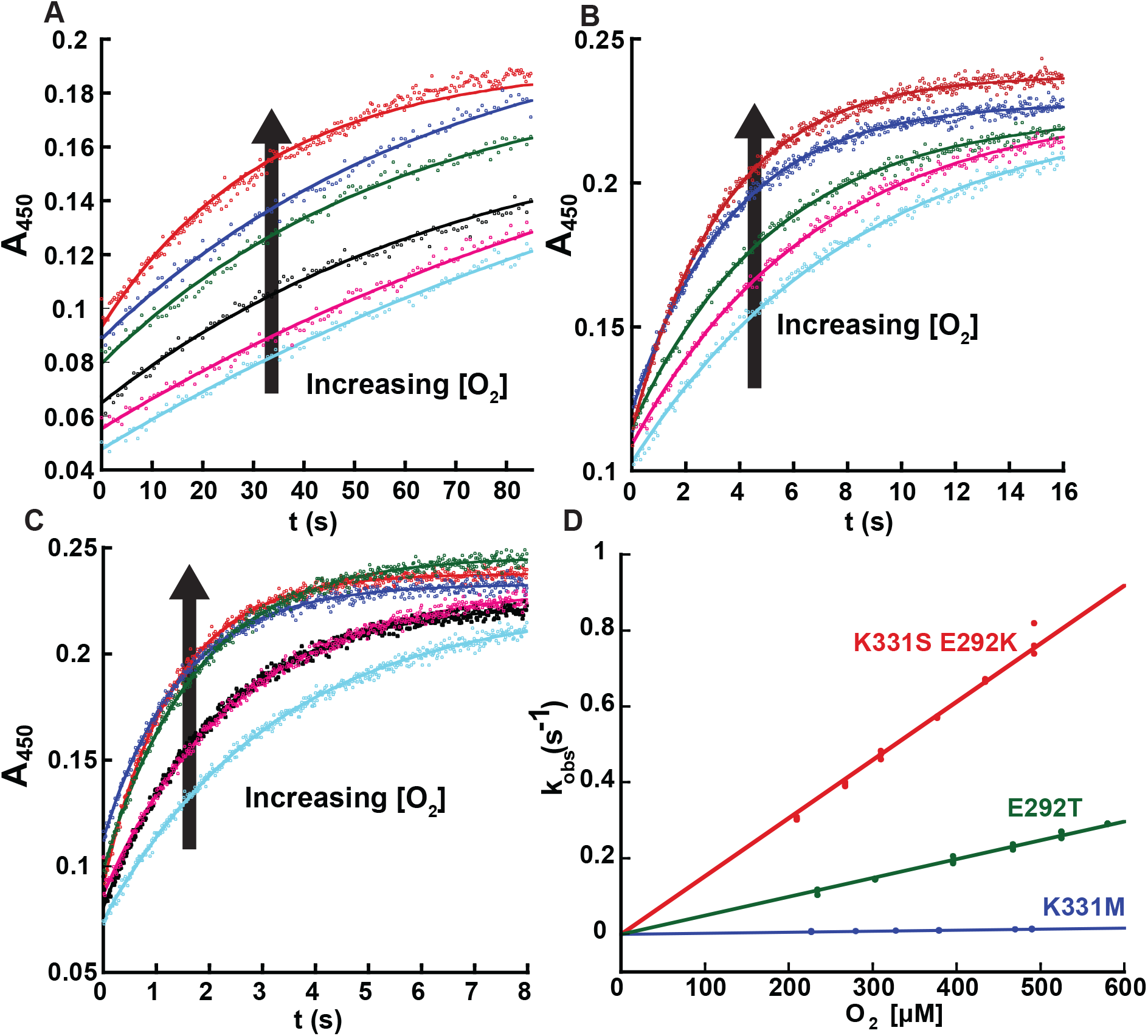
Oxidative half-reactions of NctB variants. (A), (B), and (C), Absorbance traces at 450 nm for the oxidation of K331M, E292T, and E292K/K331S variants of NctB, respectively, containing reduced flavin by various concentrations of O_2_. (D) Plots of k_obs_ values against O_2_ concentration for the reaction of various reduced NctB variants with O_2_. The slope obtained from linear fitting gives the k_ox_^O2^ values reported in Table 1.

**Fig. 5.**
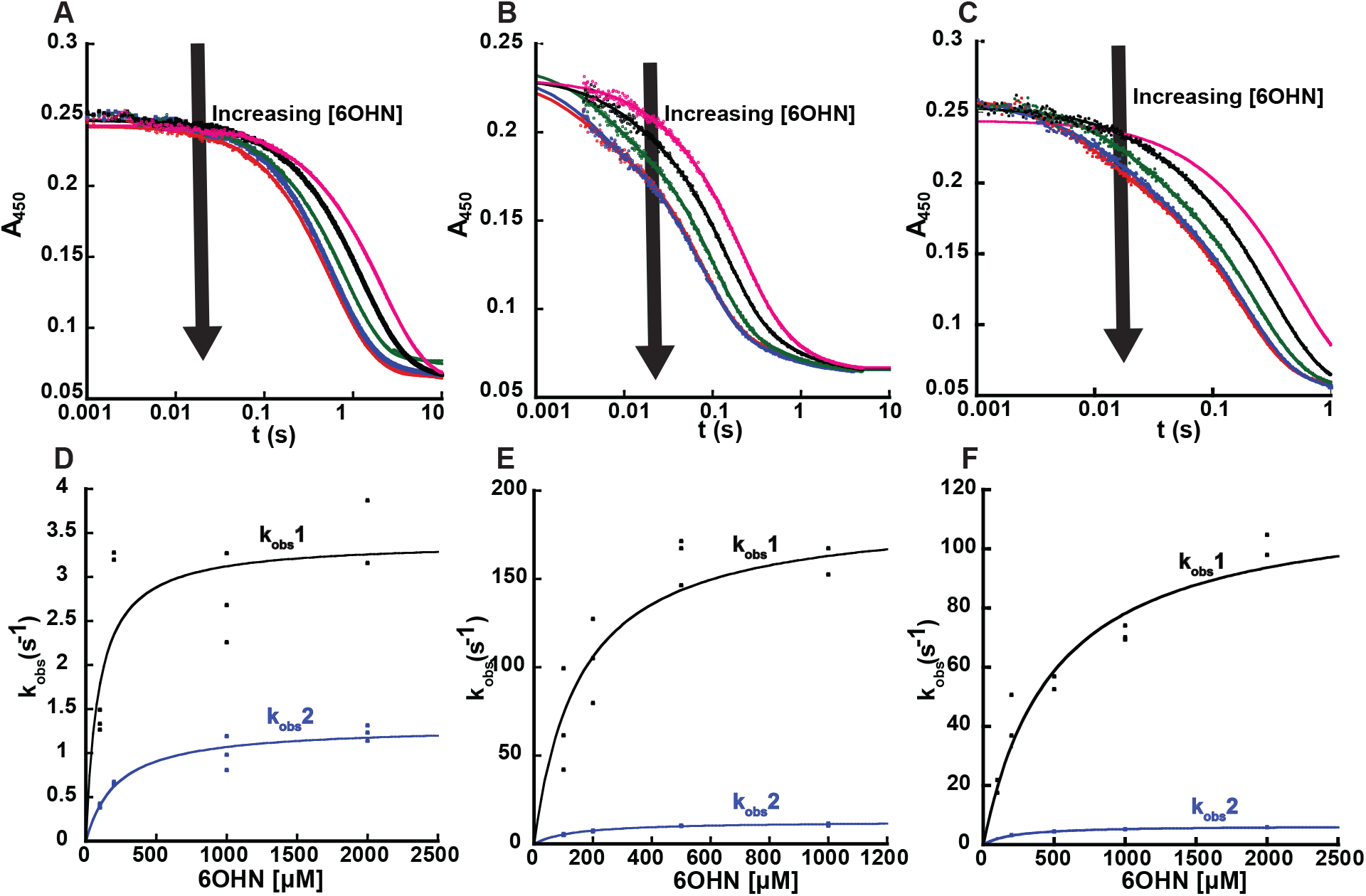
Reductive half-reactions of NctB variants. (A). (B), and (C), Absorbance trace overlay at 450 nm for the reduction of K331M, E292T, and E292K/K331S variants by various concentrations of 6OHN, respectively. Traces in (A) and (C) were fitted with equation 2 with two exponential phases. Traces in (B) for the E292T variant required a third exponential (equation 3) with a small kinetic amplitude to achieve an adequate fit. (D), (E) and (F), k_obs_ values for the first and second phase plotted against the concentration of 6OHN for NctB K331M, E292T, and E292K/K331S variants, respectively. Data for k_obs_1 is shown in black and k_obs_2 is shown in blue. Data for both k_obs_ sets are fitted to equation 4 and parameters from the resulting fit can be found in Table 1. The k_obs_ for the third, small phase for the E292T variant (not shown) was constant at 1.25 s^-1^ at all 6OHN concentrations tested and may therefore represent product dissociation.

A third variant – Glu292Lys/Lys331Ser – was constructed to install a lysine at the sequence and structural position that is present in most FAOs other than VPP-LHNO enzymes, and transient kinetic parameters for this double mutant were measured. Serine was chosen as the replacement for Lys331 because a serine is present at this position in NicA2, which is the most closely related FAO to NctB that contains a lysine at the highly conserved position (Fig. 1B and 1D). The resulting k_ox_^O2^ for the NctB Glu292Lys/Lys331Ser double mutant enzyme was 1500 M^-1^s^-1^, ∼25 fold lower compared to WT NctB (Table 1). However, the k_ox_^O2^ of this variant is ∼300 times greater than that of NicA2’s, indicating that NicA2’s lysine architecture is capable of reacting with O_2_ moderately fast if present within the appropriate enzyme context (Fig. 4C and Fig. 4D). Interestingly, the Glu292Lys/Lys331Ser variant had k_red_ values that were only ∼2-fold lower than WT (Fig. 5C, 5F and Table 1), indicating that the two mutations introduced in this variant have only a modest impact on the reductive half reaction.

### Reduction potentials

Reduction potentials of NctB and NctB variants were measured at 25°C and pH 7.5 under anaerobic conditions using the xanthine/xanthine oxidase method of Massey [27,28]. Anthraquinone-2,6-disulphonate was used as the reference dye, which has a reduction potential of -184 mV at 25°C and pH 7.5 [31]. As shown in Fig. S1 all variants and wild-type were successfully reduced in a single-stage two electron process. The linearized Nernst plots yielded a reduction potential of -193 mV for WT NctB, whereas the Lys331Met, Glu292Thr and the Glu292Lys/Lys331Ser variants had reduction potentials of -184, -203, and -202 mV respectively. Overall, the mutations to charged residues adjacent to the flavin N5 resulted in only modest changes in the reduction potential of the enzyme.

## DISCUSSION

Both NctB from *Shinella* sp. HZN7 (a VPP-LHNO enzyme) and Pyridine-LHNO from *P. nicotinovorans* catalyze the same, early step in the bacterial catabolism of nicotine by oxidizing 6OHN into 6HPON using O_2_ as the oxidant [7,24]. Both enzymes are members of the FAO superfamily, but are distantly related to each other (26% sequence identity), and phylogenetic analysis of the FAO family suggests these two enzymes independently evolved to have the same function [22,24,32,33]. Interestingly, VPP-LHNO enzymes appear to have recently evolved from within a larger clade of enzymes that function as dehydrogenases, and VPP-LHNO enzymes only recently gained the ability to react rapidly with O_2_ as an oxidant [22]. The kinetics and mechanism of Pyridine-LHNO from *P. nicotinovorans* have been extensively characterized in several studies [7,14,34]. Our kinetic results here on NctB thus allow for a direct comparison between the catalytic properties of these two enzymes that independently evolved to have LHNO function. Of particular interest is the absence of the lysine in NctB that is highly conserved among FAOs, and whether the novel lysine positioned in front of the flavin in NctB plays a similar role in O_2_ activation (Fig. 1A-C).

The results from our rapid reaction kinetic measurements demonstrate that FADH_2_ in NctB reacts rapidly with O_2_ despite having the lysine in its active site originating from a different position than that typically found in FAOs. The k_ox_^O2^ value for *P. nicotinovorans* Pyridine-LHNO has not been directly measured. However, k_cat_/K_m_ for oxygen (2.6 x 10^5^ M^-1^s^-1^), which has been measured for Pyridine-LHNO, should approximate this parameter [7]. Thus, comparing the values measured for the two enzymes (3.8 x 10^4^ M^-1^s^-1^ vs 2.6 x 10^5^ M^-1^s^-1^) shows that NctB reacts with O_2_ with a similar rate constant as Pyridine-LHNO. Notably, the measurements on Pyridine-LHNO were conducted at 25°C while our experiments on NctB were performed at 4°C, and the parameters for these two enzymes would likely be even closer if measured at the same temperature. Our measurements of the reductive half-reaction for NctB showed two kinetically observable events, with each phase contributing ∼50% of the total signal change, and the k_obs_ for both phases displaying a hyperbolic dependence on the 6OHN concentration with similar half-saturation concentrations of 6OHN. This behavior is most consistent with two populations of NctB that are reduced by 6OHN with different k_red_ values. The reaction between *P. nicotinovorans* Pyridine-LHNO and 6OHN has also been reported to occur as a biphasic process; however, in that reaction, the first phase had an amplitude ∼10 times that of the second phase, and the first phase was therefore attributed to flavin reduction while the second was attributed to release of the iminium product from the enzyme [7]. We previously observed similar biphasic flavin reduction with comparable amplitudes in the reaction of NicA2 with nicotine, which we attributed to the two protomers of the NicA2 homodimer reacting with different kinetics [12,30]. The behavior of NctB’s reductive half reaction with its substrate is therefore more similar to that of NicA2 than Pyridine-LHNO. Notably, NctB and NicA2 have a similar, extensive dimeric interface between their two protomers that is distinct from that observed in the structure of *P. nicotinovorans* Pyridine-LHNO, which may explain the biphasic flavin reduction observed in NctB and NicA2, but not in Pyridine-LHNO (Fig. S2).

The Lys331Met mutation of NctB had a dramatic effect on the kinetics of flavin oxidation by O_2_, decreasing the rate constant for this reaction ∼1400-fold (Table 1). As previously mentioned, a lysine residue that forms a water-mediated hydrogen bond with the flavin N5 is a highly conserved feature in enzymes of the FAO structural family, and mutating this residue to methionine decreases the rate constant for reaction with O_2_ in several members of the FAO family [14–21]. For example, the Lys287Met mutation decreases the k_cat_/K_m_ value for oxygen ∼6000 fold in *P. nicotinovorans* Pyridine-LHNO, highlighting the importance of this residue in promoting flavin oxidation by O_2_ in this enzyme [14]. Our results on the oxidative half reaction with the Lys331Met variant indicate that Lys331 in NctB likely plays a similar role in accelerating the reaction with O_2_ despite being in a different position than the one typically found in FAO family enzymes. The Lys331Met mutation also had a modest effect on the reductive half-reaction as well, decreasing k_red_ values 53- and 13-fold for the two observable flavin reduction events without significantly affecting the apparent K_d_ values for the substrate (Table 1). The Lys331Met mutation does not significantly alter the reduction potential of the flavin (Fig. S1), so the modest decrease in k_red_ values may instead be caused by subtle changes in positioning of the flavin cofactor upon mutating this residue, which would impact the rate constant for hydride transfer from 6OHN to the flavin N5. Prior measurements on the reductive half reaction of *P. nicotinovorans* Pyridine-LHNO showed a 11-fold decrease in k_red_ with the Lys287Met variant, which is comparable to that observed here with the Lys331Met variant of NctB [14]. Glu292 in NctB is the residue that replaces the highly conserved lysine found in most FAO enzymes (Fig. 1A). The structure of NctB shows that the side chain of Glu292 is adjacent to Lys331 (Fig. 1B), which would be expected to elevate the pKa of Lys331, and the presence of Glu292 could help stabilize the positioning of Lys331 in front of the flavin N5 to promote the reaction with O_2_. The Glu292Thr mutation decreased k_ox_^O2^ by nearly 80-fold (Table 1), demonstrating that Glu292 is indeed important for promoting a rapid reaction with O_2_, presumably due to its effect on the properties and/or positioning of Lys331. In comparison, the kinetic parameters for the reductive half reaction with the Glu292Thr variant were similar to wild type, indicating that Glu292 is not important for hydride transfer from 6OHN to the flavin.

The k_cat_/K_m_–pH profile for NctB with 6OHN was bell-shaped, with pKa values of 6.8 and 9.2. This parameter is thought to report on the protonation state of the free substrate and enzyme that are required for productive binding and catalysis [35]. The previously reported k_cat_/K_m_–pH profile for *P. nicotinovorans* Pyridine-LHNO with 6OHN was similarly bell-shaped, with pKa values of 6.2 and 9.7 [14]. The fact that the k_cat_/K_m_–pH profiles and resulting pKa values are similar for both enzymes suggests that NctB shares the same protonation state requirements as Pyridine-LHNO for the productive binding and catalysis of 6OHN. In Pyridine-LHNO, the acidic pKa was attributed in part to a hydrogen bond network of the substrate, water molecules and residues in the active site of the enzyme; this multiprotonic network gains an additional proton at low pH, introducing a positive charge, which prevents binding of 6OHN since the substrate is protonated and positively charged below its pKa value of 8.6 [14]. It is unclear if a similar hydrogen bond network exists in NctB since a structure with substrate bound has yet to be solved. However, the fact that NctB displays a similar acidic pKa in the k_cat_/K_m_–pH profile as Pyridine-LHNO suggests that a similar hydrogen bond network may exist in the 6OHN bound complex of NctB, which prevents productive 6OHN binding upon gaining a proton at low pH. The basic pKa of 9.7 in the k_cat_/K_m_–pH profile of Pyridine-LHNO with 6OHN was attributed to Lys287 in the active site of the enzyme [14]. That NctB displays a similar basic pKa of 9.2 suggests Lys331 has a similar pKa and requirement for protonation as Lys287 in Pyridine-LHNO despite Lys331 originating from a different sequence position. The k_cat_–pH profile of NctB was invariant across the pH range of 5.5-10, which is a marked departure from the bell-shaped k_cat_– pH profile of Pyridine-LHNO with pKa values of 7.0 and 10.6 [14]. The lack of a basic pKa in the k_cat_–pH profile for NctB may be due to the fact that was unstable at pH values greater than 10, which would be required to detect a pKa of ∼10.6 on k_cat_. The absence of an acidic pKa in the k_cat_–pH profile for NctB may indicate that the rate-limiting step during turnover of 6OHN is different for this enzyme than in Pyridine-LHNO.

NicA2 is an unusual member of the FAO family in that it reacts poorly with O_2_ (k_ox_^O2^ of 5 M^-1^s^-1^) in favor of using a cytochrome c as its oxidant [12,36]. NicA2 has the lysine at the highly conserved sequence position (Fig. 1A), yet reacts poorly with O_2_, suggesting that while NicA2 has the lysine known to accelerate the reaction with O_2_, other aspects of its structure restrict its ability to react with this oxidant [12,37,38]. The results on our Glu292Lys/Lys331Ser variant of NctB support this notion, as this variant reacts reasonably quickly with O_2_, with k_ox_^O2^ roughly 300-fold faster than NicA2 and 50-fold faster than the Lys331Met variant of NctB which completely lacks a lysine in its active site. This suggests that the NicA2 lysine architecture is capable of promoting the reaction of flavin hydroquinone with O_2_ if placed within a protein context that does not suppress the reaction with O_2_ (e.g., NctB). Data on the reductive half-reaction of the Glu292Lys/Lys331Ser variant shows that this variant had only a modest 2-fold reduction in k_red_ values compared to WT NctB, in contrast with the 13–50-fold reduction in k_red_ observed with the Lys331Met variant (Table 1). This result suggests that maintaining a lysine in the active site, derived from either sequence position, is necessary to react efficiently with the 6OHN substrate, which may explain why flavoprotein amine dehydrogenases like NicA2 that react poorly with O_2_ preserved the lysine in their active site despite not needing to promote the reaction with O_2_.

The results presented here examined the kinetic parameters of NctB (a VPP-LHNO), an enzyme functionally similar to yet distantly related from Pyridine-LHNO. Similar kinetic parameters, pH profiles, and mechanism of oxygen activation between both Pyridine-LHNO and NctB indicates that these two enzymes convergently evolved to have the same function. During the evolution of NctB, the enzymes gained an active site lysine at a different sequence position from that typically seen in FAOs, yet the lysine at this novel position has a similar function in accelerating the reaction with O_2_.

## DATA AVAILABILITY

All data are contained within the article. Data are available from the corresponding author on request.

## FUNDING

This research was supported by National Science Foundation grant 2236541 (to F. S.).

### CRediT author contribution statement

Zhiyao Zhang: Writing – review & editing, Writing – original draft, Visualization, Validation, Investigation, Formal analysis. Kaitrin Freeland: Investigation, Formal analysis. Frederick Stull: Writing – review & editing, Writing – original draft, Supervision, Project administration, Funding acquisition, Formal analysis, Conceptualization.

## Appendix. Supplementary Data

This article contains supporting information.

## SCHEME/FIGURE CAPTIONS

**Scheme 1.**
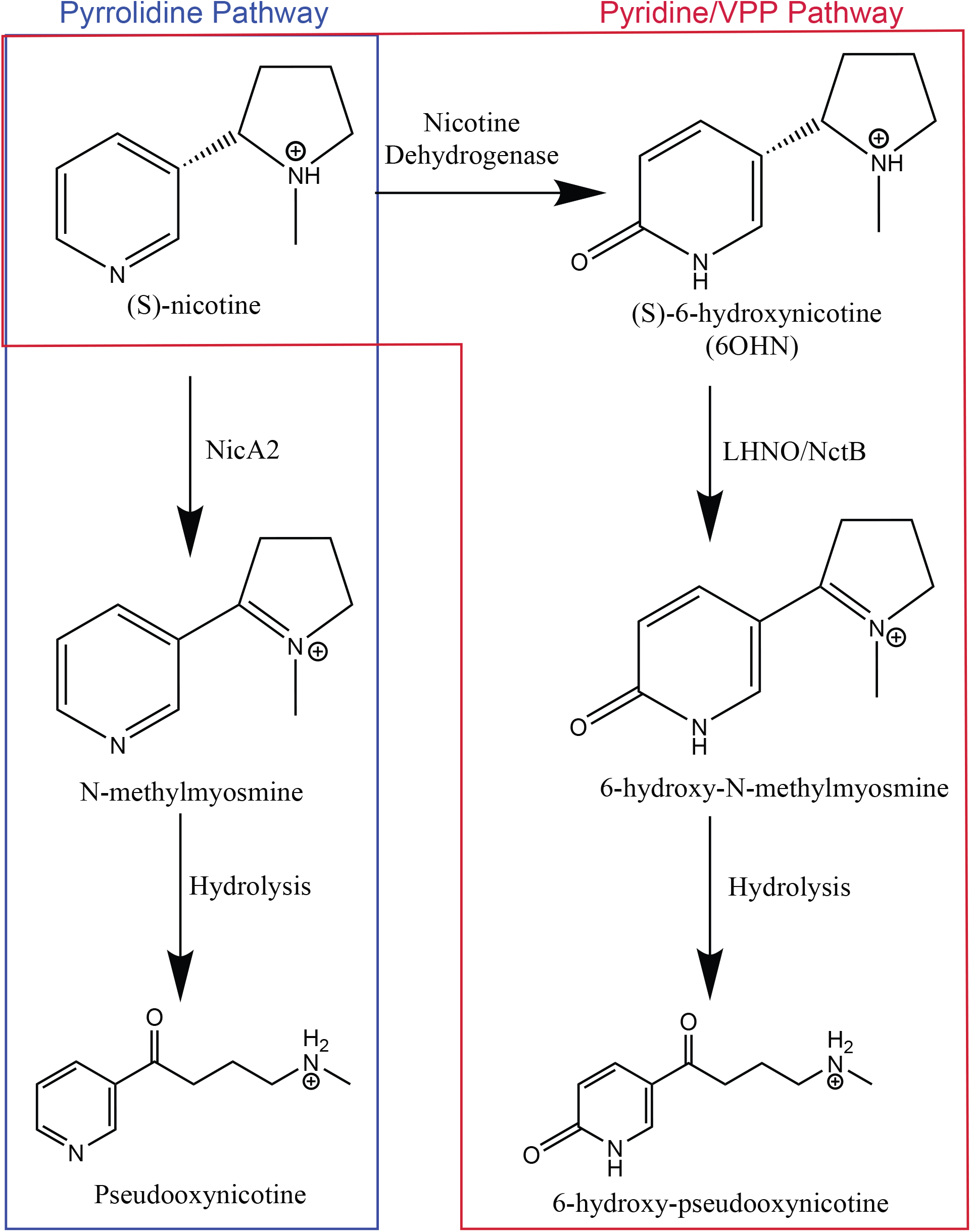
Early steps for the pyrrolidine, pyridine and VPP nicotine degrading pathways.

**Scheme 2.**
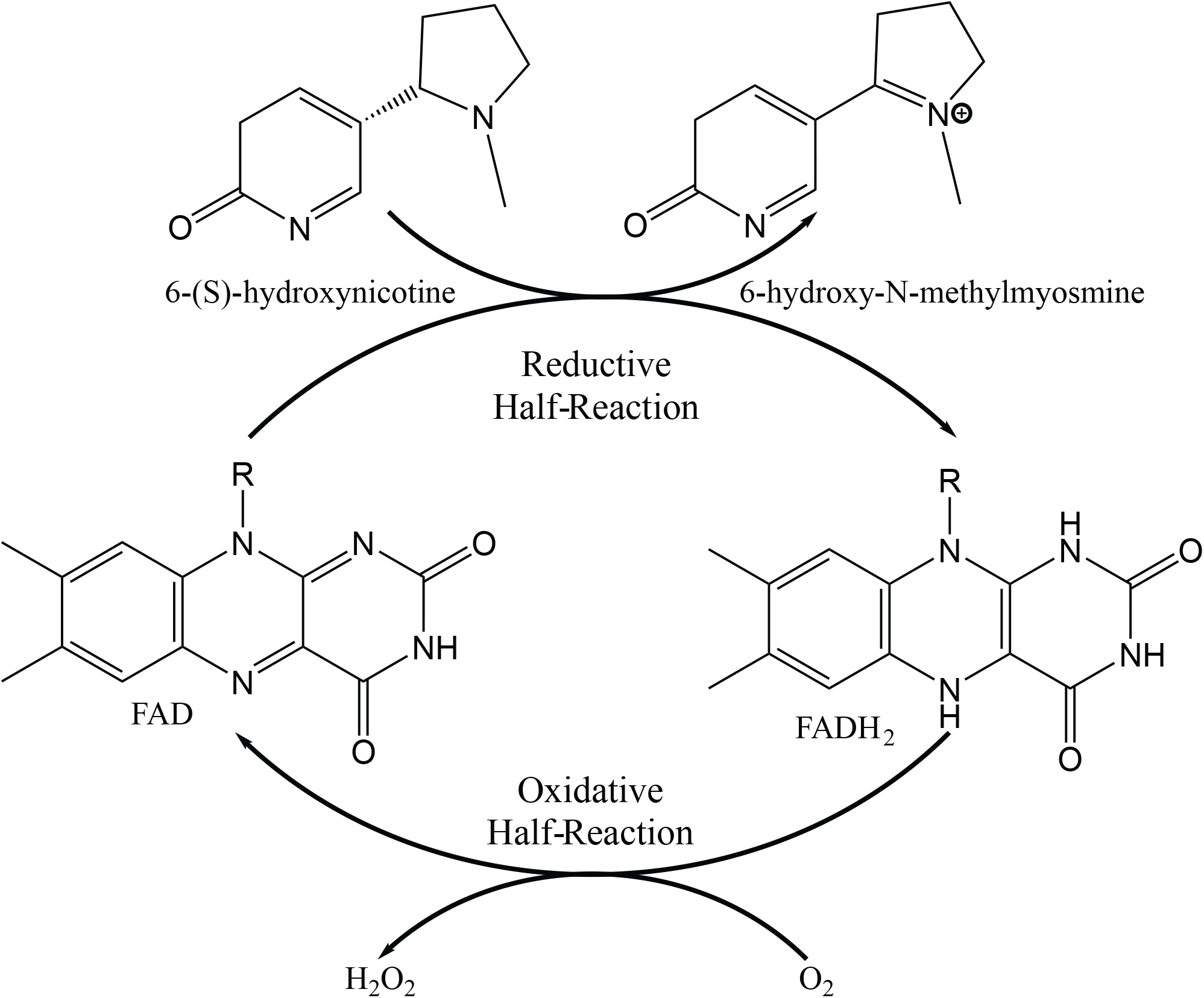
NctB catalytic cycle depicting the reductive and oxidative half-reactions.

