## Supporting information for "Role of glutamate 292 and lysine 331 in catalysis for the flavoenzyme (S)-6-hydroxynicotine oxidase from *Shinella* sp. HZN7"

### Supplementary Figures

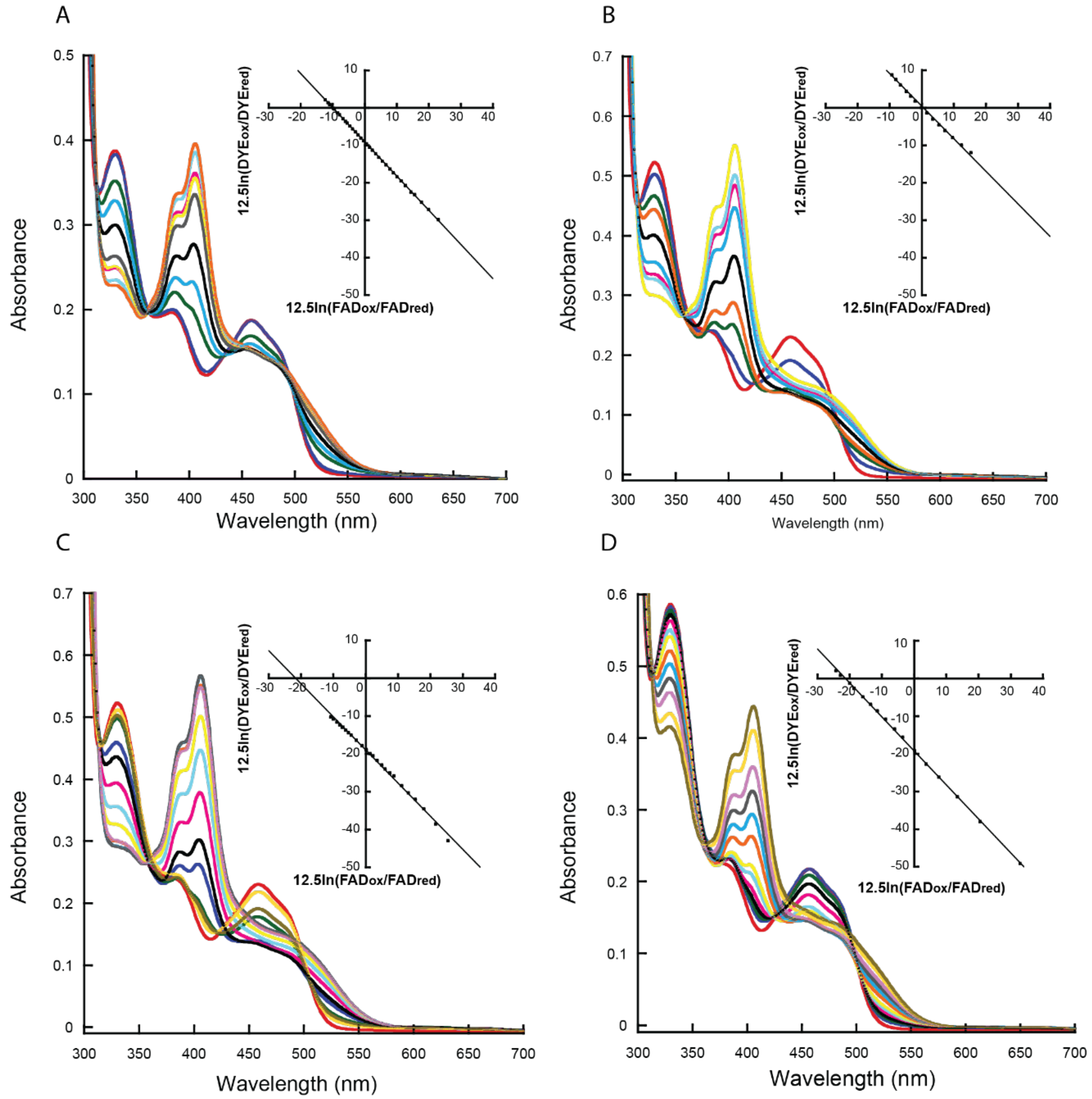


**Fig. S1. Reduction potentials of NctB and active site variants.** (A), (B), (C) and (D) Anaerobic reduction of 16 µM wild-type NctB, K331M, E292T, and E292K/K331S variants, respectively, with the xanthine/xanthine oxidase method [1,2]. Spectra were taken at time intervals (600-750s) until complete reduction in 12-18 hours. For clarity only selected spectra are shown. Insets show representative Nernst plots. The reduction of each enzyme is shown, yielding the calculated reduction potential. (A) NctB, Em =-193 mV; (B) K331M, Em = -184 mV; (C) E292T, Em = -203 mV; (D) E292K/K331S, Em = -202 mV.

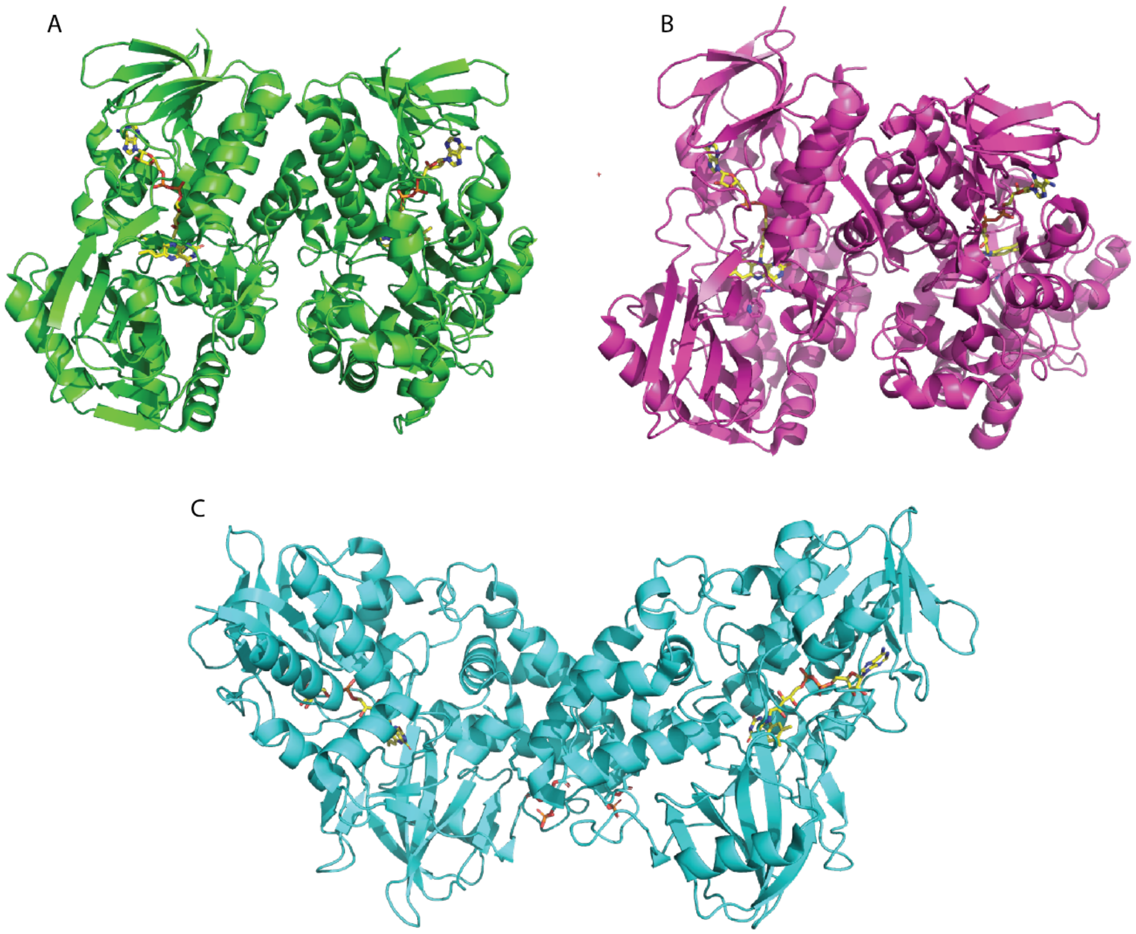


**Fig. S2. Structures of NctB, NicA2 and LHNO.** (A), (B) and (C) Crystal structures showing the dimer orientation of NctB (6CR0), NicA2 (6C71) and Pyridine-LHNO (3NG7) respectively.
